# Wheelchair user’s voice: a pilot study in Indonesia

**DOI:** 10.1101/2020.01.16.908772

**Authors:** Stephanie Gabela Vasquez, Megan E. D’Innocenzo, Jon Pearlman, Christy Zigler, Yasmin Garcia Mendez, Perth Rozen, Eviana Hapsari Dewi, Ignatius Praptoraharjo

**Affiliations:** Department of Rehabilitation Science and Technology, University of Pittsburgh, Pittsburgh, PA, United States of America; Department of Population Health Science, Duke University, Durham, NC, United States of America; United Cerebral Palsy (UCP) Wheels for Humanity, Chatsworth, California, United States of American; Department of Health Policy and Management, Gadjah Mada University, Yogyakarta, Indonesia

## Abstract

There is a significant unmet need for appropriate wheelchairs worldwide. As a whole, studies suggest that appropriate wheelchairs have a positive impact on the quality of life and health of wheelchair users, which is consistent with the goals and outcomes in more resourced settings, and that when services are provided along with the wheelchair, the positive impact is increased. The gaps in previous research, along with the global focus on evidence-based decision making, were strong motivators for carrying out a study that contrasted the outcomes associated with different types of wheelchair service provision strategies. This study used a sample of participants randomly selected from a waitlist (*N* = 142) or people who used wheelchairs as their primary means of mobility. Two different groups were included, the 8-Steps group and the Standard of Care (SOC)group. The 8-Steps group (N= 118) received wheelchairs from service providers trained using the World Health Organization (WHO) 8-Step process and the SOC group (N=24) received hospital-style wheelchairs and standard care. Interviews were conducted at baseline and a follow up 3-6 months after distribution, to collect data using the following tools: International Society of Wheelchair Professionals (ISWP) Minimum Uniform Data Set (<u>MUD</u>), Wheelchair Skills Test Questionnaire (WST-Q), and Life Satisfaction Questionnaire (LiSAT-11), and Breakdown and Adverse Consequences Questionnaire (BAC-Q).Across-group statistical comparisons were not attempted. The majority of participants from the 8-Steps group used their wheelchair every day for more than 8 hours a day. In contrast, the SOC group used their wheelchairs less than 6 hours a day. Both groups traveled less than 500 meters per day. Participants’ WST-Q scores were low, <65%, at both baseline and endline, with a significant decrease at endline. No significant differences were found when comparing device satisfaction across wheelchairs types. The majority (n=87; 72.7%) of 8-Steps group participants reported performing wheelchair maintenance. Less than half (n=9; 37.5%) of the SOC group reported performing maintenance activities. For both groups, the most reported maintenance activity was wiping or washing the wheelchair, and most wheelchair repairs were performed by the study participant or a family member. The results of this study demonstrate the importance of the WHO 8-steps training package for wheelchair provision. Further studies, training services, and wheelchair skills are needed in low and middle-income countries for both wheelchair users and service providers.

## Introduction

There is a significant unmet need for appropriate wheelchairs around the world. Using population-based estimates published by WHO, approximately 77 million people worldwide currently require the use of a wheelchair for mobility [1]. Data collected in several less-resourced settings (LRS) on access to assistive technologies suggests that only between 17% and 37% have access to appropriate assistive technologies, such as wheelchairs. Based on these data, an estimated 33-65 million people who need wheelchairs do not have access to them. This large unmet need has motivated governments, private companies, and not-for-profit organizations to provide wheelchairs through a range of largely uncoordinated service provision and supply chain approaches for the past several decades [2,3]. Concerns that some of these approaches lacked the desired impact (e.g. [4,5]) motivated a multi-year effort to establish standards related to service and product quality. A consensus conference held in 2006 led by the WHO [6] resulted in the development and publication of consensus guidelines [7] on manual wheelchair provision, and a set of consensus-based training packages to educate wheelchair service providers [8–10]. Efforts to disseminate these tools are substantial – they are widely promoted by different organizations (e.g. <u>WFOT</u>, <u>WCPT</u>, <u>ISWP</u>, <u>ISPO</u>), they are translated into several languages, and they are being adopted as the basis for global training [11,12], and competency evaluations [13].

In spite of these dissemination efforts, there has been relatively little change in the wheelchair sector, and governments, private companies, and not-for-profits continue to distribute wheelchairs that would not be considered ‘appropriate’ [6] through the service delivery approach that does not include all 8 steps recommended by WHO [7]. There are two key reasons that organizations do not universally adopt these consensus approaches. First, policies that dictate the type of wheelchair service provision are weak or non-existent in many countries where the need is greatest, and therefore organizations are not obligated to adhere to specific service or product quality standards. Second, there is a paucity of evidence that providing wheelchairs through the approach outlined by WHO, which is costlier and requires a long-term commitment, addresses the needs for wheelchairs users more efficiently or effectively.

These two reasons are closely linked and related to a lack of objective evidence about the marginal benefits of providing appropriate wheelchairs through a costlier 8-step approach (described by WHO) versus simply giving a standard hospital-style wheelchair to someone who requests it, which continues to be the standard of care in most countries. Subjective evidence indicating that hospital-style wheelchairs fail quickly in the community were published as early as 1990 [4,14], but investigated only a small number of wheelchairs and were geographically focused on India. Interest about the impact of wheelchair service increased as the sector began to coordinate in 2006 when the WHO became involved [15], and researchers began to collect and publish outcome data. For instance, a cross-sectional study on 188 wheelchair users who received basic wheelchairs without formal service revealed that 93.1% of the wheelchairs will still in use after an average of 18 months and that receiving the wheelchair was associated with a significant increase in independence and significantly decreased pressure ulcer incidence [16]. These strong positive results bolstered the argument that the costlier approach promoted by the WHO may not be necessary. Meanwhile, because the study was cross-sectional and investigated a group who received a single type of wheelchair, it does not provide conclusive evidence of the relative value of providing wheelchairs through WHO’s 8-step approach, nor provide reliable insight into whether it was the wheelchair or other factors which led the improvements. The first study we are aware of that investigated the impact of the WHO’s 8-Step service approach was in Indonesia, and compared a group receiving wheelchairs through the 8-Step process to a waitlist control group at baseline and a 6-month follow-up [17]. Subjects who received new wheelchairs reported significant increases in physical health, environmental health, and satisfaction with their mobility devices as compared to the waitlist control group. Using a robust study design and validated outcome measures, this research helps to support WHO’s 8-Step service provision approach but did not directly compare it to the standard of care. A longitudinal study of 200 individuals who received one of two designs of wheelchairs [18] was conducted in Peru, Uganda, and Vietnam found that overall health indicators, distance traveled, and employment increased, and that wheelchair design had little impact on these results. This study was conducted on a population of users similar to an earlier study [16] and similarly did not receive services based on the 8-Step approach, did not include a control group, or use strongly validated outcome measures.

The only study we are aware of that compared across service provision models was a cross-sectional study that recorded data from 852 wheelchair users in Kenya and the Philippines [19,20]. The investigators used a proxy measure for services based on the subject’s self-report of how many service steps (from 0 to 8) occurred when they received their wheelchairs. The results suggest that users in Kenya versus the Philippines were more likely to use their wheelchairs daily (60% vs. 42%) and had higher activities of daily living (ADL) performance (80% vs. 74%) highlighting the country-level differences. The impact of increased services was largely dependent on what service was received. For instance, individuals who were assessed for a wheelchair (Step 2) were more likely to have a higher ADL performance. Similarly, individuals who received training (Step 7) were more likely to use their wheelchairs daily. This cross-sectional study of a relatively large subject pool provides strong evidence of the positive impact of services on the outcomes of wheelchair service provision.

The prior research evidence paints a positive but incomplete picture of the impact of service provision in the wheelchair sector. As a whole, the studies suggest that wheelchairs have a positive impact on the quality of life and health of wheelchair users, which is consistent with the goals and outcomes in more resourced settings [21], and that the degree to which services are provided increases that impact. But there is still a significant gap in evidence related to the specific benefits of an 8-step service provision approach, versus the standard of care. This is due to limitations in the previous studies associated with the study design, such as the lack of control groups, cross-sectional methodology, or weakly validated measures. Meanwhile, the need for this information is becoming increasingly important to meet a global push towards using evidence to drive policy changes related to rehabilitation and assistive health technology purchasing decisions. These goals have been emphasized by global collaborations such as through Call to Action in WHO’s REHAB2030 [23], WHO’s GATE Research Priorities [23], and AT scale [25].

The gaps in previous research along with the global focus on evidence-based decision making motivated our team to carry out a study that contrasted the outcomes associated with different types of wheelchair service provision strategies. This study design was tailored to identify hypotheses of potential outcomes and inform changes to, a wheelchair supplier (Consolidating Logistics for Assistive Technology Supply & Provision), whose goal is to sell a range of appropriate wheelchair models to buyers who then provide them through a global service network. The study was guided by the following research questions:

1. What are the key challenges to performing a longitudinal controlled wheelchair study in Indonesia?
2. What are some of the hypotheses related to the wheelchair model and its potential effects on key outcome variables such as performance, usability, reliability, and quality of life in Indonesia?

## Methods

A longitudinal, mixed-methods study was carried out to evaluate the impact of wheelchair service provision from three wheelchair providers (WPs) in Indonesia: Puspadi, the Bunga Bali Foundation (BBF), and the Social Department. Puspadi is staffed by service providers who were all trained to provide services using the 8-Step service provision model described in the WHO guidelines [7], whereas BBF and Social Department used the standard-of-care where they distributed hospital-style wheelchairs to those who requested them without any clinical services.

Three research teams were involved in the project and secured IRB approval. A team from the Comprehensive Initiative on Technology Evaluation (CITE) at the Massachusetts Institute of Technology designed the initial study and supported in-country data collection but was not involved in the final selection of the Standard of Care group. A team from Center for Health Policy and Management (<u>CHPM</u>), Gadjah Mada University led the data collection efforts in Indonesia. A team from the Department of Rehabilitation Sciences and Technology (<u>RST</u>) from the University of Pittsburgh led data analysis and drafting of this manuscript. The study was supported with a grant from Google.org (grant #322068) which was awarded to United Cerebral Palsy - Wheels for Humanity (<u>UCP-W</u>) who contracted the other organizations to carry out the research. IRB approval was secured at Gadjah Mada University, RST, and MIT.

Wheelchair users on the waitlist from Puspadi and BBF and the Social Department were recruited into the study. Users who were 16 years or older, could interact and communicate help caregiver help, were recruited to participate in the study. The target sample size was limited by the size of the waitlist, which was just over 200 people. The sample size was also limited by the number of wheelchairs that were available at Puspadi, BBF, the Social Department as well as the study costs for wheelchairs.

Wheelchair users receiving wheelchairs from Puspadi were provided with one of five different wheelchair models according to their needs: Standard (Std.), Motivation Active Folding (MAF), UCP Expression (UCP), Rough Rider (RR), and Motivation Rough Terrain (MRT). These wheelchairs are shown in Figure 1. Puspadi wheelchair service providers had received training using WHO packages to provide wheelchairs according to the 8-Step approach. Wheelchair users receiving wheelchairs from BBF were given a basic hospital-style wheelchair (H), see Figure 1. Individuals providing wheelchairs at BBF had not been formally trained.

**Figure 1.**
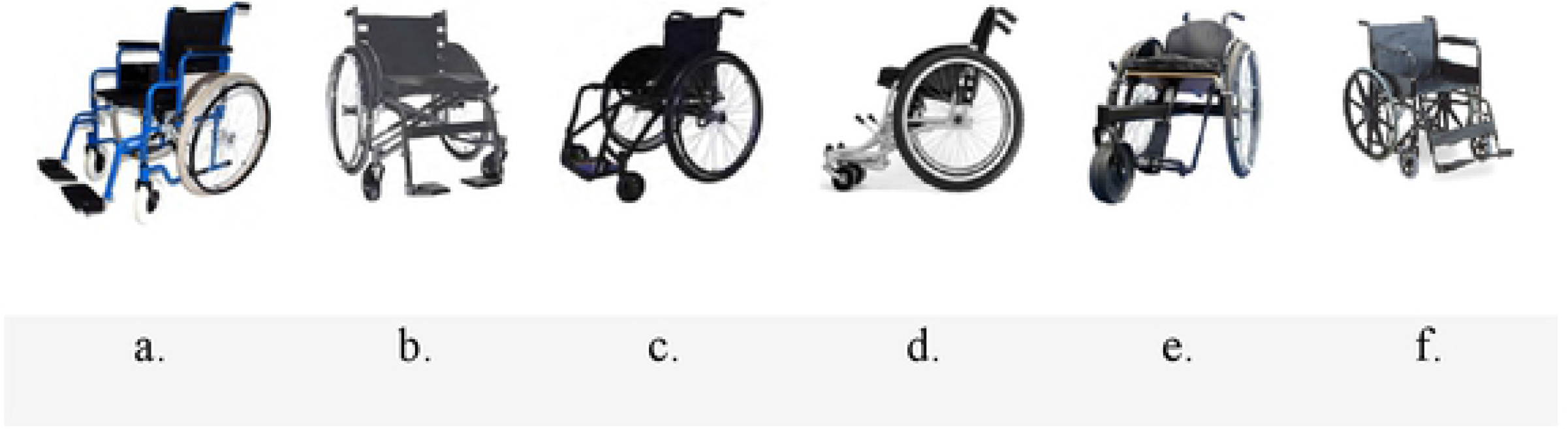
Types of wheelchairs provided, a. Std, b. MAF, c. UCP, d. RR, e. MRT, f. H.

## Data collection methods & tools

Data was collected from the subjects through in-person interviews that were used to record responses to a set of standardized questionnaires. The close-ended responses were entered into tablets by researchers from the CHPM Gadjah Mada University team using KoboToolbox, a survey software. The interviews were recorded, and the responses to open-ended questions were transcribed and translated into English by the research team at Gadjah Mada University.

The interview protocol was comprised of questions from a variety of questionnaires including the International Society of Wheelchair Professionals Minimum Uniform Data Set (ISWP-<u>MUD</u>), Wheelchair Skills Test Questionnaire (WST-Q) [25], Poverty Probability Index (PPI) [26] for Indonesia, and Life Satisfaction Questionnaire (LiSAT-11) [27], Breakdown and Adverse Consequences Questionnaire (BAC-Q) [29], Quebec User Satisfaction with Assistive Technology (QUEST) and Functional Mobility Assessment (FMA). We adapted all of the questionnaires to the local context in several ways. First, they were all were translated into one of the two the local Indonesian languages, Balinese or Bahasa. Second, the tools were modified to fit the cultural context based on testing in the field and feedback from local partners, such as the UCP Roda Untuk Kemanusiaan, Puspadi, BBF, and CHPM Gadjah Mada University. The questionnaires were tested with similar types of wheelchair users before the data collection period.

Table 1 provides an overview of the types of data collected through each questionnaire. These questionnaires were administered at baseline and endline, 3-6 months after the start of the study, to all the wheelchair users who participated in the study.

**Table 1:** Questionnaires Administered at Baseline and Endline.

| Tool | Data Type |
| --- | --- |
| ISWP-MUD | Demographics, wheelchair usage (self-report) |
| PPI* | Likelihood of being under the poverty line |
| WST-Q | Wheelchair skills |
| BAC | Wheelchair repairs |
| LiSAT-11 | Life satisfaction |
| QUEST | Perception of satisfaction with assistive technology and services |
| FMA | Self-reported effectiveness of wheeled mobility devices |

\* data not reported at endline due to missing and incomplete data

## Data analysis & statistical evaluation

Baseline demographic characteristics were compared between the 8-Steps group and Standard of Care group using independent samples t-tests and chi-squared tests (for continuous and categorical variables, respectively) to ensure comparable groups. Self-reported wheelchair usage, in terms of days per week, hours per day, distance traveled and the places where the device was used were reported for each group, and the change from baseline to endline was described. The McNemar tests were used to determine significant consistency in the reported settings where wheelchairs were used. Additionally, stacked columns were used to compare frequency count across types of wheelchairs for wheelchair usage using Microsoft Excel.

Significant changes in wheelchair skills (as per WST-Q total scores) were also evaluated for the 8-Steps and the Standard of Care groups independently. The WST-Q was recorded for participants who owned a manual wheelchair at the beginning of the study, and to the participants who had used the wheelchair provided by the end of the study. Missing responses were considered as not valid scores. As owning a wheelchair would be important to the amount of change that we would expect, an independent samples t-test was used to determine differences in WST-Q scores between people who owned and did not own a wheelchair at baseline. A paired samples t-test within each group was used to compare changes between baseline and endline for those participants who owned a manual wheelchair at baseline and used the wheelchair provided during the study. Graphical analysis was performed across wheelchairs as part of secondary analysis for WST-Q. Wheelchair maintenance and repairs were analyzed using frequency statistics. IBM SPSS Statistics for Windows Version 25.0 (IBM Corp.) was used to perform all statistical analyses (alpha = .05).

ISWP-MUD gathered information about the participant’s satisfaction with the study wheelchair. This information was analyzed using the Wilcoxon-signed rank test to determine changes from baseline to endline. As a secondary/exploratory analysis, a Kruskal-Wallis test was used to determine differences in satisfaction for people receiving different wheelchair models. Life satisfaction was assessed at baseline and endline using the LiSat-11 questionnaire, which was comprised of 11 items, concerning life as a whole, vocation, economy, leisure, contacts, sexual life, ADL, family life, partner, physical health, and psychological health. The question about sexual health was removed due to the sensitive nature of the question, leaving 10 questions. Satisfaction was estimated across a six-level scale (from 1 = very dissatisfied to 6 = very satisfied), higher scores indicating higher levels of life satisfaction. A total score can be calculated (range: 10–60) [32].

## Results

A total of 150 participants were recruited for the study, 15% of whom had not owned wheelchairs previously. A total of eight participants were excluded from data analysis, six that did not participate in the follow-up, and two that were deceased before the conclusion of the study. Therefore, longitudinal data from 142 participants was analyzed; 118 from the 8-Steps groups and 24 from the Standard of Care group.

Descriptive statistics of age, gender, mobility aid use, disability, and education level are shown for each group in Table 2. There were no significant differences between the groups for gender (p=.169). However, individuals in the SOC group were significantly older (p=.001) and were less likely than the 8-Steps group to report using a mobility aid at enrollment (p=.001; Table 2). There were also differences in reported diagnoses between groups. More than half of the participants recruited from the 8-Steps group participants had polio (51.7%), but no participants from the SOC group reported having polio. Due to the important differences between these two groups, subsequent results are presented separately.

**Table 2:**
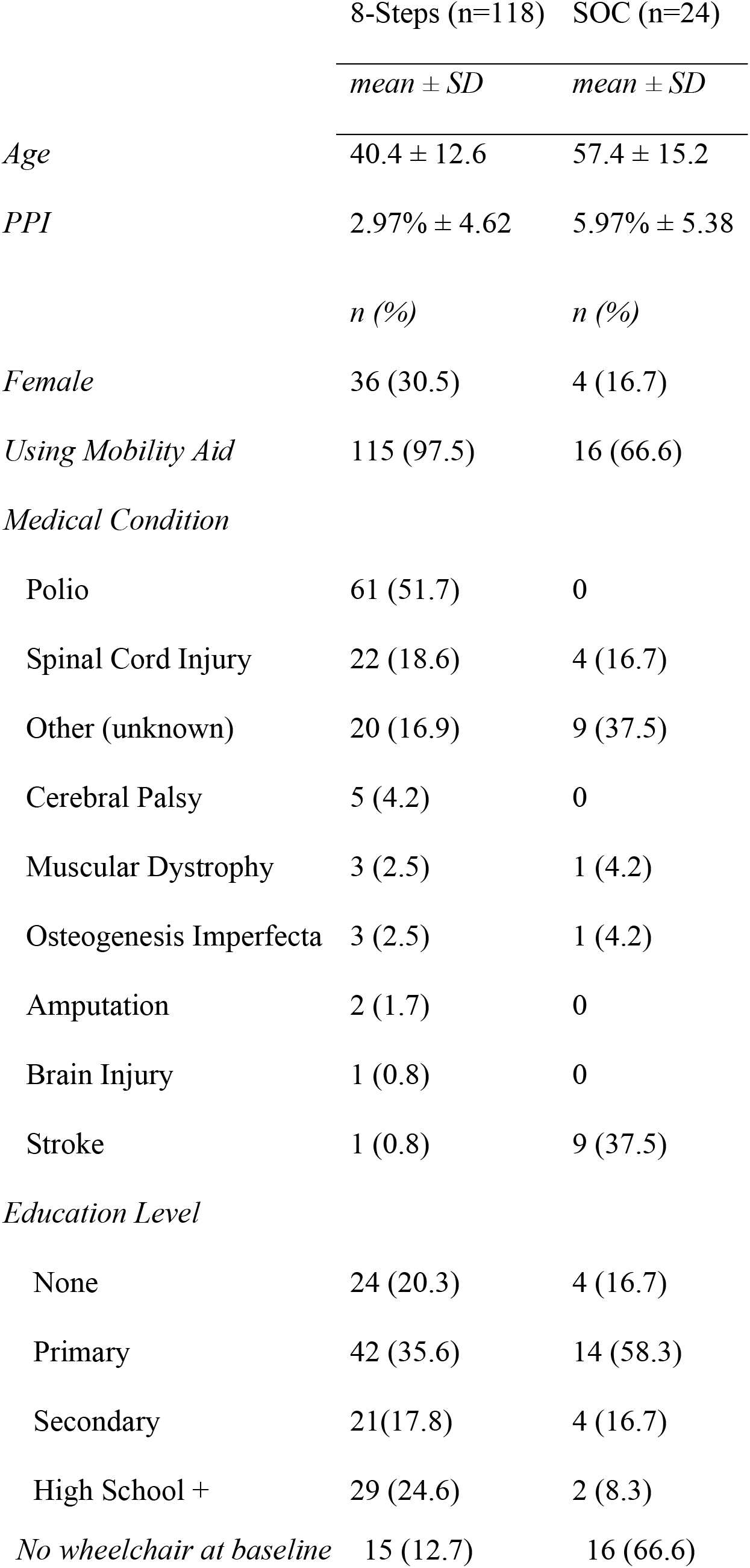
Demographics.

The majority of the participants from8-Steps group used their wheelchair every day and traveled <500 meters per day at baseline and endline (Table 3). There were no major changes in the self-reported wheelchair usage between baseline and endline (Table 3). For usage measured by ‘days per week,’ the increase in the number of individuals reporting using their wheelchair 1-3 days a week at endline was almost entirely attributed to the individuals who did not have a wheelchair at baseline. Wheelchair usage was similar for individuals in the SOC group (Table 3); the majority of participants used the device every day and traveled <500m. However, the majority of individuals in this group reported using their wheelchair between 1-6 hrs./day. More information about change over time for wheelchair usage (in terms of days per week and distance traveled per day) can be found in the Appendix.

**Table 3:**
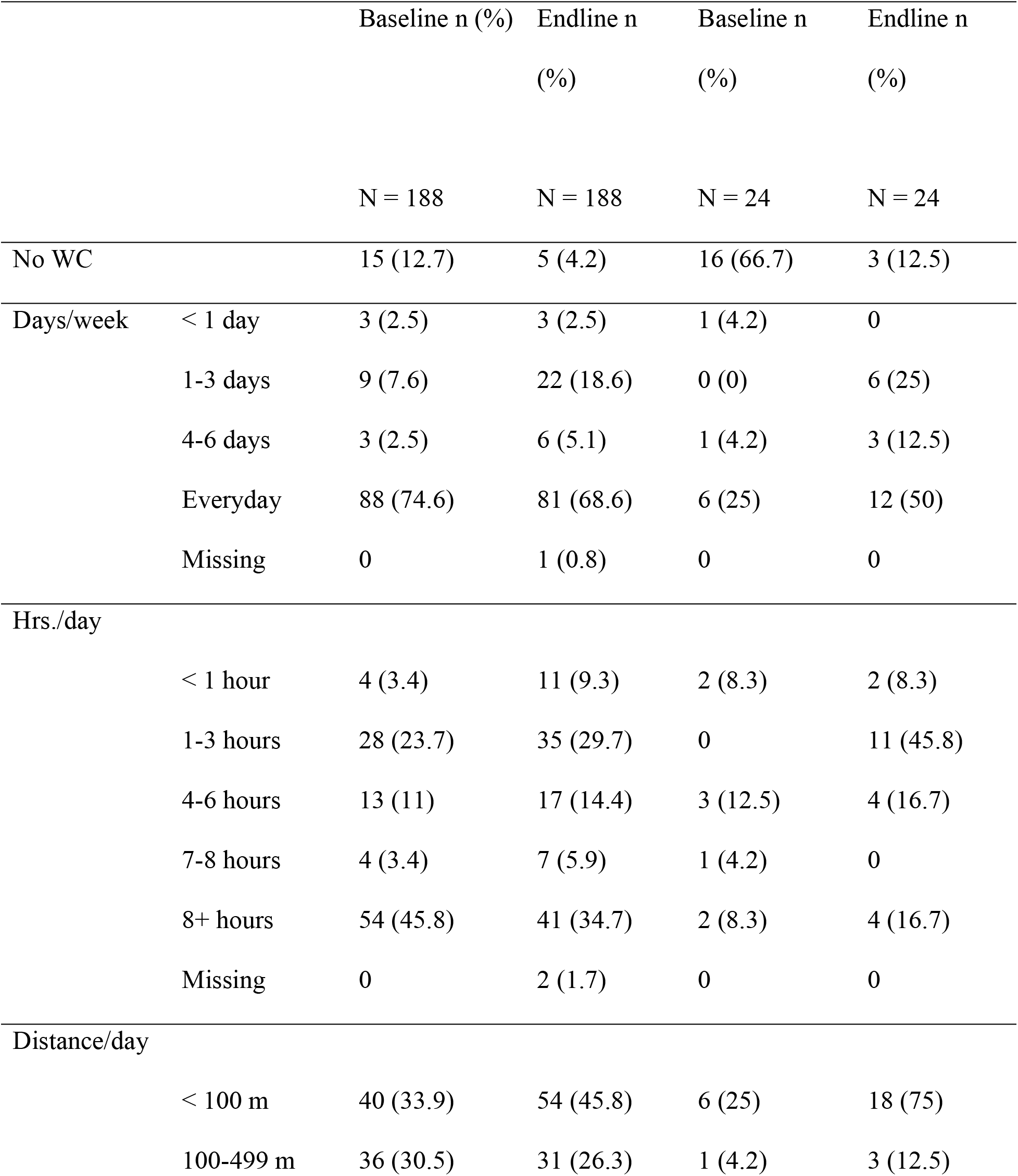
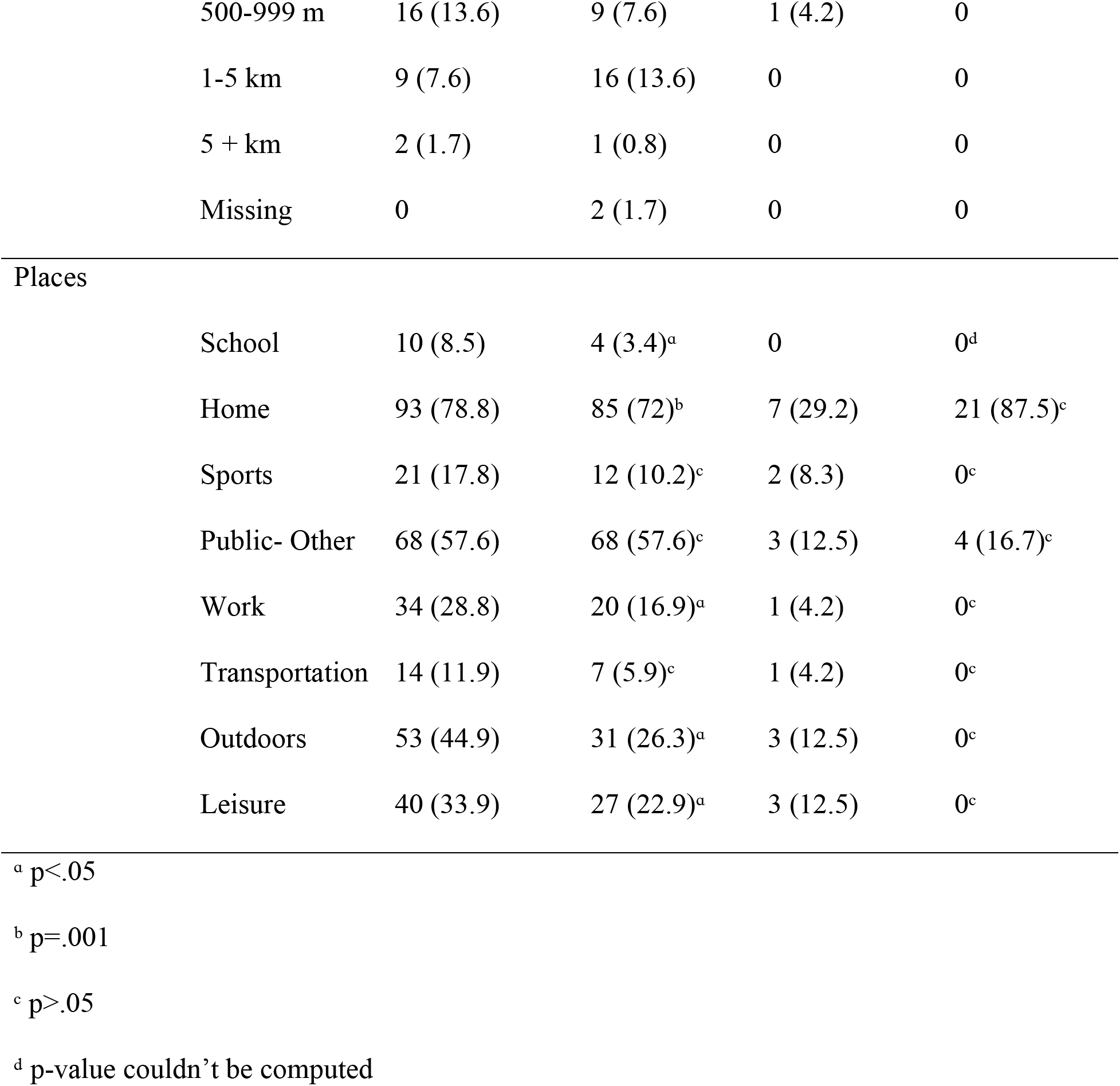
wheelchair usage descriptive statistics.

|  |  | Baseline n (%) | Endline n<br>(%) | Baseline n<br>(%) | Endline n<br>(%) |
| --- | --- | --- | --- | --- | --- |
|  |  | N = 188 | N = 188 | N = 24 | N = 24 |
| No WC |  | 15 (12.7) | 5 (4.2) | 16 (66.7) | 3 (12.5) |
| Days/week | < 1 day | 3 (2.5) | 3 (2.5) | 1 (4.2) | 0 |
|  | 1-3 days | 9 (7.6) | 22 (18.6) | 0 (0) | 6 (25) |
|  | 4-6 days | 3 (2.5) | 6 (5.1) | 1 (4.2) | 3 (12.5) |
|  | Everyday | 88 (74.6) | 81 (68.6) | 6 (25) | 12 (50) |
|  | Missing | 0 | 1 (0.8) | 0 | 0 |
| Hrs./day |  |  |  |  |  |
|  | < 1 hour | 4 (3.4) | 11 (9.3) | 2 (8.3) | 2 (8.3) |
|  | 1-3 hours | 28 (23.7) | 35 (29.7) | 0 | 11 (45.8) |
|  | 4-6 hours | 13 (11) | 17 (14.4) | 3 (12.5) | 4 (16.7) |
|  | 7-8 hours | 4 (3.4) | 7 (5.9) | 1 (4.2) | 0 |
|  | 8+ hours | 54 (45.8) | 41 (34.7) | 2 (8.3) | 4 (16.7) |
|  | Missing | 0 | 2 (1.7) | 0 | 0 |
| Distance/day |  |  |  |  |  |
|  | < 100 m | 40 (33.9) | 54 (45.8) | 6 (25) | 18 (75) |
|  | 100-499 m | 36 (30.5) | 31 (26.3) | 1 (4.2) | 3 (12.5) |
|  | 500-999 m | 16 (13.6) | 9 (7.6) | 1 (4.2) | 0 |
|  | 1-5 km | 9 (7.6) | 16 (13.6) | 0 | 0 |
|  | 5 + km | 2 (1.7) | 1 (0.8) | 0 | 0 |
|  | Missing | 0 | 2 (1.7) | 0 | 0 |
| Places |  |  |  |  |  |
|  | School | 10 (8.5) | 4 (3.4)<sup>a</sup> | 0 | 0<sup>d</sup> |
|  | Home | 93 (78.8) | 85 (72)<sup>b</sup> | 7 (29.2) | 21 (87.5)<sup>c</sup> |
|  | Sports | 21 (17.8) | 12 (10.2)<sup>c</sup> | 2 (8.3) | 0<sup>c</sup> |
|  | Public- Other | 68 (57.6) | 68 (57.6)<sup>c</sup> | 3 (12.5) | 4 (16.7)<sup>c</sup> |
|  | Work | 34 (28.8) | 20 (16.9)<sup>a</sup> | 1 (4.2) | 0<sup>c</sup> |
|  | Transportation | 14 (11.9) | 7 (5.9)<sup>c</sup> | 1 (4.2) | 0<sup>c</sup> |
|  | Outdoors | 53 (44.9) | 31 (26.3)<sup>a</sup> | 3 (12.5) | 0<sup>c</sup> |
|  | Leisure | 40 (33.9) | 27 (22.9)<sup>a</sup> | 3 (12.5) | 0<sup>c</sup> |

<sup>a</sup> p<.05
<sup>b</sup> p=.001
<sup>c</sup> p>.05
<sup>d</sup> p-value couldn't be computed

The participants from the 8-Steps group reported usage in all settings at both time points (Table 3). The most frequently reported setting was ‘home’, but individuals also commonly used their wheelchair in ‘other public places’ and ‘outdoors on rough surfaces.’ The 8-Steps group had significant differences in almost all settings (Table 3). For example, 23 individuals at baseline who reported using their wheelchair at ‘home,’ reported not using their wheelchair at ‘home’ at endline. Similar negative changes were seen in other settings like ‘school,’ ‘work,’ ‘outdoors on rough surfaces,’ and ‘leisure activities.’ The only two settings reported by this group at endline were ‘home’ and ‘other public places’. The SOC group had very small cell sizes, and thus, the differences were not statistically significant. At endline, 5(4.2%) participants from the 8-Steps group and 3 (12.5%) individuals from the SOC group reported they were not using the study wheelchair.

Overall, most of the individuals receiving any of the wheelchairs reported using it every day, except for individuals who received the MRT (Figure S5). Participants who received an MRT were more likely to report using it only 1-3 days per week although a number of them still reported using it every day. Interestingly, usage in hours per day was bimodal; individuals were most likely to report using their wheelchair 1-3 hours per day or 8+ hours per day (Figure S5).

Individuals who received RR or Std were more likely to report using it the most (8+ hours per day), while individuals who received H or UCP were slightly more likely to report lower usage (1-3 hours per day). Additionally, the majority of participants reported traveling less than 100 m per day (Figure S5). This was similar for all types of wheelchairs in the 8-Steps group. Individuals who received H wheelchairs reported traveling the shortest distance, with no one having this type of wheelchair with a distance traveled over 500m. Individuals with RR and MRTs seemed to travel the longest distances, although some individuals with Std, UCP, and MAF’s did report traveling >1km.

It is important to note that individuals could choose multiple settings where they used their wheelchair. Although wheelchair use at ‘home’ was the most frequently reported setting for all wheelchair types (Figure 2), there were some interesting differences in the other settings by wheelchair type. The participants who used H only reported usage at ‘home’ and ‘other public places.’ In contrast, those participants who used RR tended to report more settings and choose those settings that were less frequently chosen by the sample such as ‘sports,’ ‘work,’ and ‘transportation.’ UCP users also reported usage in all settings but not as high as RR users.

**Figure 2.**
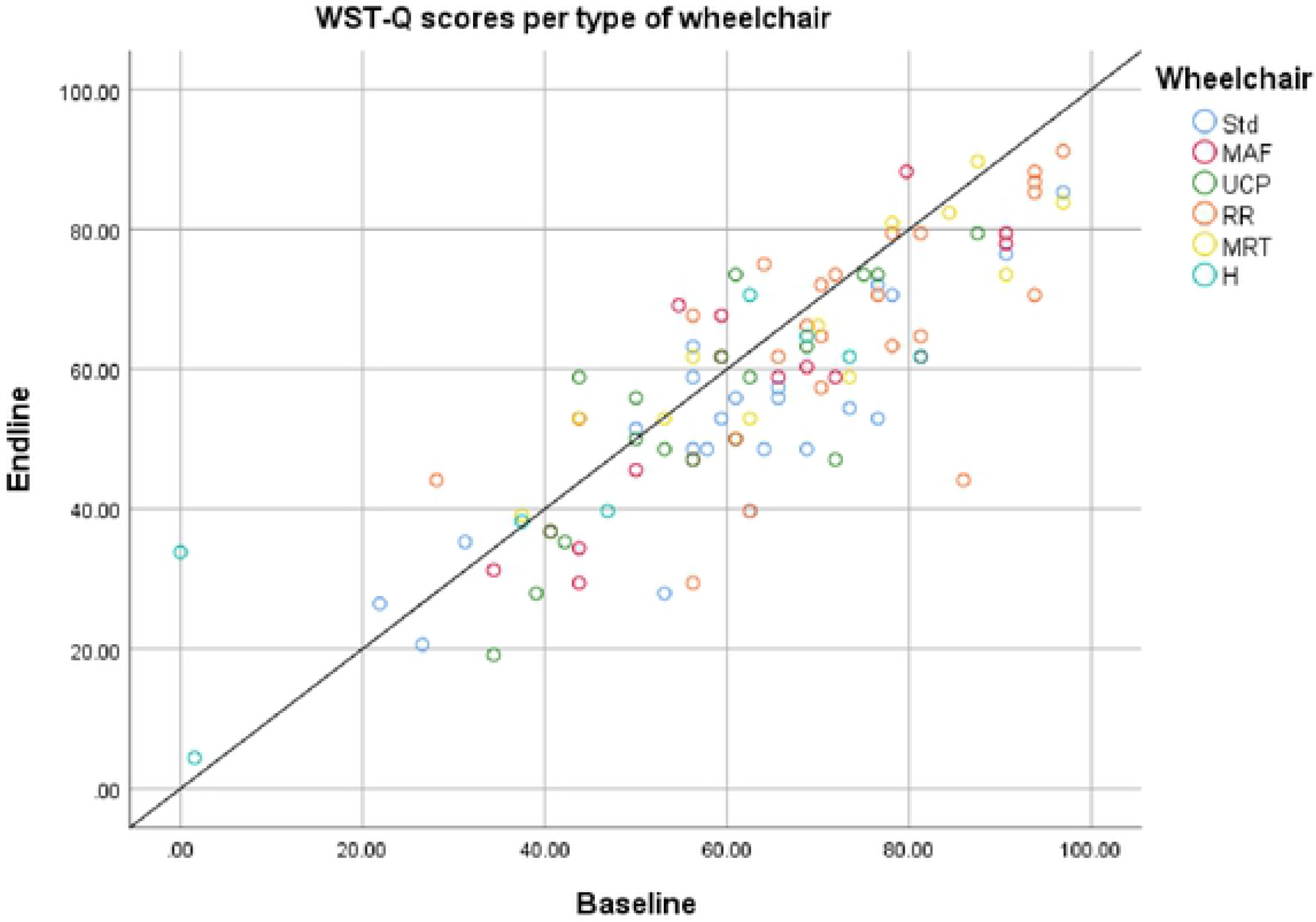
WST-Q scores per type of wheelchair.

## Wheelchair skills

In the group, 15 (12.7%) participants did not own a wheelchair at baseline and did not have baseline WST-Q scores. At endline, the 8-Steps group had 8 (6.9%) participants with non-valid scores. From the 5 (4.2%) individuals who did not use the wheelchair from the study, 2 (1.7%) of them also reported not owning a wheelchair at baseline, therefore, the tool was not administered. Thus, the final sample used for analysis was 92. Overall, average total WST-Q scores decreased from baseline (M=64.7, SD=17.9) to endline (M=58.6, SD=17.2) with an average decrease of 6.03 points (SD=10.4). This difference was statistically significant, t (91) = 5.542, p<.001, d=.577. There were no significant differences in final wheelchair skills for individuals who owned a wheelchair at baseline compared to those who did not own one. An independent sample t-test was conducted to determine whether there was a difference in WST-Q scores at endline between those participants who owned a wheelchair at baseline (n=103) and those did not own one (n=15). At endline, participants without a wheelchair at baseline (M=59.926, SD=16.539) had almost identical average skills when compared to those who owned a wheelchair at baseline (M=58.619, SD=17.218), t (103) =.257, p=.797, d=.076.

In the Standard of Care group, the majority of participants (16/24; 66.6%) did not own a wheelchair at baseline and 3 (12.5%) of them did not use the wheelchair provided. Also, one of the participants obtained a score of zero at baseline as all the individual skills were responded as “no” when asked about the capacity of doing them. Due to the extremely low sample size at each time-point, statistical tests for wheelchair skills were not performed in this group. In terms of wheelchair skills at baseline, For the 8 individuals with scores at baseline, the mean WST-Q score was 46.48 (SD=31.50). Regarding wheelchair skills for the 21 individuals with scores at endline, the mean WST-Q score was 34.31 ± 25.29.

Individuals in the 8-Steps group reported more basic skills and more intermediate skills than individuals in the SOC group (Table 4). In other words, more than 85% of participants from the 8-Steps group reported being able to roll short distances, turn, maneuver sideways, transfer on and to level surfaces, and roll on soft surfaces. On the other hand, less than 63% of the participants from the SOC group reported mastery of these skills, see Table 4. However, a small number of participants from both groups reported capacity in advanced skills like ascending steep inclines and performing wheelies.

**Table 4:** individual skills for WST-Q capacity at endline.

| Skill Level | Individual Skill | 8-Steps | SOC |
| --- | --- | --- | --- |
|  |  | *n*(%) | *n*(%) |
| Basic | Rolls forward short distance | 108 (91.5) | 12 (50) |
| Basic | Rolls backward short distance | 103 (87.3) | 13 (54.2) |
| Basic | Turns in place | 103 (87.3) | 15 (62.5) |
| Basic | Turns while moving forward | 111 (94.1) | 16 (66.7) |
| Basic | Turns while moving backward | 106 (89.8) | 13 (54.2) |
| Basic | Maneuvers sideways | 105 (89) | 10 (41.7) |
| Basic | Reaches high object | 64 (54.2) | 6 (25) |
| Basic | Pick object from floor | 96 (81.4) | 13 (54.2) |
| Basic | Operate body positioning options | 88 (74.6) | 12 (50) |
| Basic | Relieves weight from buttocks | 99 (83.9) | 13 (54.2) |
| Basic | Level transfer | 101 (85.6) | 14 (54.2) |
| Intermediate | Folds and unfolds wheelchair | 71 (60.2) | 7 (29.2) |
| Intermediate | Gets through hinged door | 90 (76.3) | 8 (33.3) |
| Intermediate | Rolls longer distance | 87 (73.7) | 12 (50) |
| Intermediate | Avoids moving obstacles | 91 (77.1) | 8 (33.3) |
| Intermediate | Ascends slight incline | 89 (75.4) | 5 (20.8) |
| Intermediate | Descends slight incline | 98 (81.3) | 8 (33.3) |
| Intermediate | Rolls across side-slope | 64 (54.2) | 7 (29.2) |
| Intermediate | Rolls on soft surface | 102 (86.4) | 14 (58.3) |
| Intermediate | Gets over threshold | 89 (75.4) | 7 (29.2) |
| Intermediate | Gets over gap | 43 (36.4) | 5 (20.8) |
| Intermediate | Ascends low curb | 46 (39) | 4 (16.7) |
| Intermediate | Descends low curb | 53 (44.9) | 8 (33.3) |
| Advanced | Ascends steep incline | 18 (15.3) | 0 |
| Advanced | Descends steep incline | 35 (29.7) | 0 |
| Advanced | Ascends high curb | 11 (9.3) | 1 (4.2) |
| Advanced | Descend high curb | 24 (20.3) | 2 (8.3) |
| Advanced | Performs stationary wheelie | 29 (24.6) | 3 (12.5) |
| Advanced | Turns in place in wheelie position | 19 (16.1) | 2 (8.3) |
| Advanced | Descends high curb in wheelie position | 11 (9.3) | 0 |
| Advanced | Descends steep incline in wheelie position | 13 (11) | 0 |
| Advanced | Gets from the ground into wheelchair | 79 (66.9) | 10 (41.7) |
| Advanced | Ascends stairs | 8 (6.8) | 2 (8.3) |
| Advanced | Descends stairs | 3 (2.5) | 0 |

On a secondary analysis by wheelchair type, RR users reported higher wheelchair skills scores (half >70%) when compared to individuals with H wheelchairs (the majority having <50%). Over time, highly skilled MAF and MRT users showed an increase in their WST-Q scores at the end of the study while H users showed only a small increase compared to baseline (Figure 2).

## Device satisfaction

As part of the ISWP-MUD, the participants were asked to rate the satisfaction with their wheelchair from 1 (not satisfied) to 5 (very satisfied). The 8-Steps group had a rate of M=4.06, SD= 1.04 at baseline and M=4.15, SD=.99 at endline. The BBF group was slightly less satisfied overall at baseline (M=3.88, SD=.83), but on average, it increased (M=4.28, SD=.64) at endline. No significant differences were found for the participants’ satisfaction rate about the device between both time points. Table 5 shows the satisfaction rate per type of wheelchair. No significant differences were found when compared across wheelchairs.

**Table 5:** satisfaction rate based on the type of wheelchair.

|  | Not satisfied |  |  |  | Very Satisfied |  |
| --- | --- | --- | --- | --- | --- | --- |
|  | 1 | 2 | 3 | 4 | 5 | Missing |
|  | 1 (3.8) | 1 (3.8) | 2 (7.7) | 7 (26.9) | 12 (46.2) | 2 (7.7) * |
| Std. | 0 (0) | 2 (8.7) | 2 (8.7) | 9 (39.1) | 8 (34.8) | 0 (0) * |
| MAF | 0 (0) | 2 (9.1) | 2 (9.1) | 9 (40.9) | 9 (40.9) | 0 (0) |
| UCP | 0 (0) | 1 (3.8) | 3 (11.5) | 7 (26.9) | 14 (53.8) | 1 (3.8) |
| RR | 0 (0) | 0 (0) | 5 (23.8) | 6 (28.6) | 7 (33.3) | 1 (4.8)* |
| MRT | 0 (0) | 0 (0) | 2 (8.3) | 11 (45.8) | 8 (33.3) | 0 (0)* |
| H | 1 (3.8) | 1 (3.8) | 2 (7.7) | 7 (26.9) | 12 (46.2) | 2 (7.7) * |

\*The totals do not add to 100% as some participants did not use the wheelchair from the study (Std=1, MAF=2, MRT=2, H=3).

**Table 6:** LiSAT-11 8-Steps and SOC groups self-reported satisfaction at baseline and endline.

|  | 8-Steps (n=118) |  | SOC (n=24) |  |
| --- | --- | --- | --- | --- |
|  | *Baseline* | *Endline* | *Baseline* | *Endline* |
| Life as a whole | 4.1 (1.4) | 4.2 (1.5) | 3.9 (1.2) | 4.3 (.97) |
| Vocation | 3.2 (2.2) | 3.6 (2.1) | 3.7 (1.4) | 3.3 (1.3) |
| Economy | 3.3 (1.6) | 3.3 (1.7) | 3.6 (1.4) | 2.7 (1.3) |
| Leisure | 4.0 (1.5) | 3.9 (1.7) | 4.2 (1.2) | 4.8 (.43) |
| Contact | 4.9 (.95) | 4.7 (1.2) | 4.3 (1.4) | 4.7 (.77) |
| Activities of Daily Living | 4.8 (1.0) | 4.7 (1.2) | 4.3 (1.2) | 4.5 (.96) |
| Family Life * | 4.5 (1.2) | 4.6 (1.2) | 4.7 (1.1) | 4.3 (1.1) |
| Partner Relationship ** | 5.1 (.80) | 4.8 (.83) | 5.1 (.07) | 4.7 (.91) |
| Physical Health | 4.2 (1.3) | 4.0 (1.6) | 4.2 (1.3) | 3.9 (1.5) |
| Psychological Health | 4.5 (1.2) | 4.3 (1.6) | 4.7 (1.0) | 4.0 (1.4) |

\* participants living with $\geq 1$ family member 8-Steps (n=110) SOC (n=23)
\*\* participants who reported having a partner 8-Steps (n=55) SOC (n=16)

## Wheelchair maintenance and repair

The 8-Steps group reported 34 (28.8%) subjects had wheelchairs that stopped functioning correctly or broke. The most common complaint was one or more parking brakes no longer functioned properly 9 (7.6%), followed by a bearing stopped turning smoothly 8 (6.8%). Some other wheelchair repairs included tire replacement, broken wheels, and tire inflation. Of those repairs recorded, 12 (10.2%) were performed by the participant or a family member followed by the service that provided the wheelchair 11 (9.3%). Two individuals (1.7%) in the SOC group had wheelchairs that stopped functioning correctly or had a broken wheel. In both instances, the participant or a family member performed the repair.

The majority (n=87; 72.7%) of participants reported performing wheelchair maintenance. The most-reported maintenance activity was wiping or washing the wheelchair 54 (45.8%) followed by adding oil 20 (16.9%) and adding air to the tires 11 (9.3%). 8-Steps group or family members did most of the wheelchair repairs 79 (66.9%). A total of 9 subjects from the SOC group reported performing maintenance activities, 4(16.7%) subjects reported wiping or washing the wheelchair followed by 2(8.3%) added air to the tires. All participants that reported wheelchair maintenance, mentioned it was performed by the participant or a family member.

Both the 8-Steps group and SOC group reported increased satisfaction in life as a whole. The 8-Steps group also reported increased satisfaction in vocation and family life, while the SOC group reported increased satisfaction in leisure, contact, and activities of daily living.

## Discussion

The present study describes the characteristics of wheelchair usage, skills, maintenance and repairs, and life satisfaction for individuals who received wheelchair services in the 8-Steps group and those who did not receive services but still received wheelchairs (SOC group). The fact that the majority of participants owned a wheelchair when they were recruited in the study could be the reason why the patterns of wheelchair usage were not considerably different at the end line. In other words, most of them used the new wheelchair in a similar way they did with their previous one. The adherence to the new wheelchair was high, as more than 95% from the 8-Steps group and more than 87% of participants from the SOC group, were still using the study wheelchair at endline. Although not directly comparable, users who received their wheelchair through the 8-step process from Puspadi had more usage daily, hourly, reported more distance traveled per day and more places where the device was used than the BBF group. This could be due to several factors, such as the provision of a wheelchair that did not meet their needs, it did not fit properly to their body, lack of user and maintenance training, environmental barriers, or due to differences in the population. These findings are aligned with those of the study published by Toro et al. [17] in Indonesia that suggested the positive impact of the WHO 8-steps in wheelchair provision.

The total scores for wheelchair skills were overall low for participants from both groups, however, the lowest scores were obtained by the participants who received wheelchairs from BBF and had no previous wheelchair. The average scores obtained in this study at endline (8-Steps group (≥ 70 %) SOC group (≤ 50 %), were considerably lower compared to the scores to previous studies. For example, the study done by Hosseini et al. obtained an average score of 84% in the wheelchair skills test [29]. The average total score in the study by Kirby et al. [31] on the questionnaire version was 84.8%. In the study done by Toro et al. [17] in Indonesia, the participants who received wheelchair services with the WHO 8-Steps had scores of 70.6% for adults and 77.7% for adults and proxy. One reason why our results are different from the studies by Hosseini et al [29] and Kirby et al. [30] might be because all of the participants from these studies had spinal cord injury, relied exclusively on a manual wheelchair for mobility, and therefore might have had more training and practice. Our study population included subjects with a diverse set of disabilities, with Polio being most common (Table 2). The study by Toro et al. [17] had similar conditions as our study in terms of the application of the WHO 8-Steps and also happened in Indonesia. However, their findings determined no significant differences in wheelchair skills between baseline and endline.

In contrast, in this study, wheelchair skills *decreased significantly* for those who owned a wheelchair at the beginning of the study and received services following the WHO 8-steps. This could imply that they found their new wheelchair more difficult to maneuver, in part because they had not fully adjusted to it. These results suggest additional wheelchair skills training may be necessary when a wheelchair user receives a replacement or a new wheelchair. As Toro et al. [17] emphasized, the wheelchair skills included in the WHO WSTP only include seven basic skills, that consist of pushing, turning, moving up and down slopes, moving up and down steps with assistance, and doing a partial wheelie. Our results suggested that almost all of the participants who had service providers following the WHO guidelines were able to push and turn. However, more training may be needed relating to moving up and down slopes and holding a wheelie. These skills are necessary for different environments indoors and outdoors as not being able to perform them can affect their community participation, self-esteem, quality of life, work and school attendance, and more. Learning these skills allows people to become more independent as maneuvering their wheelchair and having control over it is important to perform different activities in multiple settings. Even though there is an overall positive impact in following the WHO guidelines for wheelchair provision [17], it is necessary to provide education more about mobility skills to service providers so that they can provide training about more mobility skills to wheelchair users. This should, in fact, be considered not only in the WHO WSTP packages but also in educational programs as mentioned by Fung et al. [31] Besides, educational and training programs should include information not only about mobility and skills, but also about maintenance and repairs to improve wheelchair provision and services around the world.

Future studies that consider WHO 8-steps as guidelines for wheelchair service provision could consider including training in more mobility skills. To measure the impact of both parameters in low- and middle-income countries, other studies could also consider children, youth, adults, and elderly people who have an appropriate wheelchair and are trained in multiple wheelchair skills. Further, additional studies could analyze the environment and accessibility conditions to be able to measure the impact of both appropriate wheelchair provision and an accessible environment so that social participation, work, school attendance, could be measured more objectively in developing countries. Overall, it would be useful to see more rigorous studies carried out in less-resourced countries to continue to improve wheelchair provision.

## Limitations

The biggest limitation of the study was that the subject groups were not randomly assigned to wheelchair groups. Individuals from the 8-Steps group and the SOC group were significantly different; therefore, explicit group comparisons were not attempted. During the project, it seemed that at times there were not enough resources to fully complete the 8-step process and it was challenging to determine how well a group of providers is adhering to the 8-step process. In addition, some of the wheelchair users received a wheelchair three to four months before the baseline due to low distribution resources. This created issues with recall and confusion about which wheelchair type the data collectors were referring to in the study questions.

There are a variety of outcomes and impacts that could result from users having access to proper wheelchairs, training, and services. Some outcomes and impacts of having and using a wheelchair do not appear within a three- to six-month period, making it hard to measure all of the outcomes in this study.

One of the biggest limitations of this study was the difficulty of collecting data from participants from both groups at baseline and endline interviews. Even though all questionnaires were translated into local languages and responses were translated into English, the reliability of the translated questionnaires is still unknown. A reason for the missing data could be that some questions were not clear to participants. The lack of responses made it difficult to calculate and evaluate scores from validated questionnaires. Data loss and missing data implies challenges with data collection and may have led to questionable or biased results. In future studies, it will be important to limit the length of questionnaires to avoid participant fatigue, confirm that participants fully understand all of the questions, and ensure that the questionnaires are tested for reliability.

## Conclusion

Our results provide general support that wheelchair users who are provided wheelchairs by service providers trained in the WHO 8-Step process have positive outcomes. We also found that outcomes are impacted by the wheelchair model used, reinforcing the need for proper assessments and a range of available wheelchairs. Our results support the need for increased wheelchair skills training to ensure that users learn how to use their new wheelchairs, and also can safely navigate through their environment. Finally, our study highlights many of the challenges of performing outcomes research in this population and environment that should be taken into consideration when designing robust research studies in less-resourced environments.

## Acknowledgements

The authors would like to thank the personnel of UCP Wheels for Humanity, Center for Health Policy and Management (CHPM), Gadjah Mada University,–the Puspadi Bali, the Comprehensive Initiative on Technology Evaluation (CITE) at the Massachusetts Institute of Technology, specifically Kendra Leith and Julia Heyman, Dan Frey, and Nancy Adams.

## Supporting information

S1. Fig

***S2. Fig***

|  | No WC | < 100<br>m | 100 m -<br>499 m | 500 m -<br>999 m | 1-5 km | > 5 km | Missing | **Total** |
| --- | --- | --- | --- | --- | --- | --- | --- | --- |
| No WC | 2 | 8 | 2 | 2 | 1 | 0 | 0 | 15 |
| < 100 m | 1 | 25 | 8 | 2 | 2 | 0 | 2 | 40 |
| 100 m-499 m | 2 | 16 | 11 | 4 | 3 | 0 | 0 | 36 |
| 500 m - 999 m | 0 | 4 | 4 | 0 | 7 | 1 | 0 | 16 |
| 1-5 km | 0 | 1 | 4 | 1 | 3 | 0 | 0 | 9 |
| > 5 km | 0 | 0 | 2 | 0 | 0 | 0 | 0 | 2 |
| **Total** | 5 | 54 | 31 | 9 | 16 | 1 | 2 | 118 |

***S3. Fig***

***S4. Fig***.

## References

1. Joint Position Paper on the Provision of Mobility Devices in Less-resourced Settings: A Step Towards Implementation of the Convention on the Rights of Persons with Disabilities (CRPD) Related to Personal Mobility. World Health Organization; 2011.

2. Pearlman J, Cooper RA, Zipfel E, Cooper R, McCartney M. Towards the development of an effective technology transfer model of wheelchairs to developing countries. Disabil Rehabil Assist Technol. 2006;1: 103–110.

3. Pearlman J, Cooper R, Krizack M, Lindsley A, Wu Y, Reisinger K, et al. Technical and Clinical Needs for Successful Transfer and Uptake of Lower-Limb Prosthetics and Wheelchairs in Low Income Countries. IEEE-EMBS Magazine (In Press). 2007;

4. Mukherjee G, Samanta A. Wheelchair charity: a useless benevolence in community-based rehabilitation. Disabil Rehabil. 2005;27: 591–596.

5. Kim J, Mulholland S. Seating/Wheelchair Technology in The Developing World: Need for A Closer Look. Technol Disabil. 1999;11: 21–27.

6. Sheldon S, Jacobs NA. Report of a Consensus Conference on Wheelchairs for Developing Countries: Bengaluru, India 6-11 November 2006. ISPO; 2007.

7. World Health Organization, WHO. Guidelines on the Provision of Manual Wheelchairs in Less Resourced Settings. World Health Organization; 2008.

8. WHO | Wheelchair Service Training Package - Basic level. World Health Organization; 2017; Available: http://www.who.int/disabilities/technology/wheelchairpackage/en/

9. WHO | Launching of WHO Wheelchair Service Training Package for Managers and Stakeholders. World Health Organization; 2016; Available: http://www.who.int/disabilities/technology/wheelchairpackage/wstpmanagers/en/

10. Organization WH, Others. Wheelchair service training of trainers package. World Health Organization; 2017; Available: http://apps.who.int/iris/bitstream/10665/258701/4/9789241512398-managers-eng.pdf

11. Burrola-Mendez Y, Goldberg M, Gartz R, Pearlman J. Development of a Hybrid Course on Wheelchair Service Provision for clinicians in international contexts. PLoS One. journals.plos.org; 2018;13: e0199251.

12. Burrola-Méndez Y, Goldberg M. Development and Implementation of a Hybrid Wheelchair Workshop for Clinicians in International Settings. wheelchairnetwork.org. Available: http://wheelchairnetwork.org/wp-content/uploads/2018/01/ISS2018_HybridWorkshopPaper_20180220_YB.docx

13. Gartz R, Goldberg M, Miles A, Cooper R, Pearlman J, Schmeler M, et al. Development of a contextually appropriate, reliable and valid basic Wheelchair Service Provision Test. Disabil Rehabil Assist Technol. 2017;12: 333–340.

14. Saha R, Dey A, Hatoj M, Poddar S. Study of Wheelcahir Operations in Rural Areas Covered Under the District Rehabilitaton Centre (DRC) Scheme. Indian Journal of Disability and Rehabilitation. 1990; 74–87.

15. Sheldon S, Jacobs NA. ISPO consensus conference on wheelchairs for developing countries: Conclusions and recommendations. Prosthet Orthot Int. Taylor & Francis; 2007;31: 217–223.

16. Shore SL. Use of an economical wheelchair in India and Peru: impact on health and function. Med Sci Monit. 2008;14.

17. Toro ML, Eke C, Pearlman J. The impact of the World Health Organization 8-steps in wheelchair service provision in wheelchair users in a less resourced setting: a cohort study in Indonesia. BMC Health Serv Res. 2016;16: 26.

18. Shore S. The long-term impact of wheelchair delivery on the lives of people with disabilities in three countries of the world. Afr J Disabil. 2017;6: 344.

19. Bazant ES, Himelfarb Hurwitz EJ, Onguti BN, Williams EK, Noon JH, Xavier CA, et al. Wheelchair services and use outcomes: A cross-sectional survey in Kenya and the Philippines. Afr J Disabil. 2017;6: 318.

20. Williams E, Hurwitz E, Obaga I, Onguti B, Rivera A, Sy TRL, et al. Perspectives of basic wheelchair users on improving their access to wheelchair services in Kenya and Philippines: a qualitative study. BMC Int Health Hum Rights. 2017;17: 22.

21. Chaves ES, Boninger ML, Cooper R, Fitzgerald SG, Gray DB, Cooper RA. Assessing the influence of wheelchair technology on perception of participation in spinal cord injury1. Arch Phys Med Rehabil. Elsevier; 2004;85: 1854–1858.

22. Rehab 2030 Meeting Report [Internet]. World Health Organization; Available: https://www.who.int/disabilities/care/Rehab2030MeetingReport2.pdf?ua=1

23. Global priority research agenda for improving access to high-quality affordable assistive technology [Internet]. World Health Organization; Available: http://apps.who.int/medicinedocs/documents/s23346en/s23346en.pdf

24. ATscale Global Strategy [Internet]. AT Scale; Available: https://static1.squarespace.com/static/5b3f6ff1710699a7ebb64495/t/5b55d6fb1ae6cf630bb0be94/1532352251993/Final+ATscale_2pager.pdf

25. Mountain AD, Kirby RL, Smith C. The wheelchair skills test, version 2.4: Validity of an algorithm-based questionnaire version. Arch Phys Med Rehabil. 2004;85: 416–423.

26. Desiere S, Vellema W, D’Haese M. A validity assessment of the Progress out of Poverty Index (PPI)^TM^. Eval Program Plann. Elsevier; 2015;49: 10–18.

27. Post MW, van Leeuwen CM, van Koppenhagen CF, de Groot S. Validity of the Life Satisfaction questions, the Life Satisfaction Questionnaire, and the Satisfaction with Life Scale in persons with spinal cord injury. Arch Phys Med Rehabil. 2012;93: 1832–1837.

28. Toro ML, Worobey L, Boninger ML, Cooper RA, Pearlman J. Type and Frequency of Reported Wheelchair Repairs and Related Adverse Consequences Among People with Spinal Cord Injury. Arch Phys Med Rehabil. 2016;97: 1753–1760.

29. Hosseini SM, Oyster ML, Kirby RL, Harrington AL, Boninger ML. Manual wheelchair skills capacity predicts quality of life and community integration in persons with spinal cord injury. Arch Phys Med Rehabil. 2012;93: 2237–2243.

30. Kirby RL, Worobey LA, Cowan R, Pedersen JP, Heinemann AW, Dyson-Hudson TA, et al. Wheelchair Skills Capacity and Performance of Manual Wheelchair Users with Spinal Cord Injury. Arch Phys Med Rehabil. 2016;97: 1761–1769.

31. Fung KH, Rushton PW, Gartz R, Goldberg M, Toro ML, Seymour N, et al. Wheelchair service provision education in academia. Afr J Disabil. 2017;6: 340.

32. Melin, R., Fugl-Meyer, K. S., & Fugl-Meyer, A. R. (2003). Life satisfaction in 18- to 64- year-old swedes: In relation to education, employment situation, health and physical activity. Journal of Rehabilitation Medicine, 35(2), 84–90. doi:10.1080/165019703104075597

33. Kirby, R.L., Smith, C., Parker, K., McAllister, M., Boyce, J., Rushton, P.W., … Brandt, A. (2016) The Wheelchair Skills Program Manual version 4.3 “low tech, high impact”. Retrieved from http://www.wheelchairskillsprogram.ca.

